# Structural polymorphism and diversity of human segmental duplications

**DOI:** 10.1101/2024.06.04.597452

**Authors:** Hyeonsoo Jeong, Philip C. Dishuck, DongAhn Yoo, William T. Harvey, Katherine M. Munson, Alexandra P. Lewis, Jennifer Kordosky, Gage H. Garcia, Human Genome Structural Variation Consortium (HGSVC), Feyza Yilmaz, Pille Hallast, Charles Lee, Tomi Pastinen, Evan E. Eichler

## Abstract

Segmental duplications (SDs) contribute significantly to human disease, evolution, and diversity yet have been difficult to resolve at the sequence level. We present a population genetics survey of SDs by analyzing 170 human genome assemblies where the majority of SDs are fully resolved using long-read sequence assembly. Excluding the acrocentric short arms, we identify 173.2 Mbp of duplicated sequence (47.4 Mbp not present in the telomere-to-telomere reference) distinguishing fixed from structurally polymorphic events. We find that intrachromosomal SDs are among the most variable with rare events mapping near their progenitor sequences. African genomes harbor significantly more intrachromosomal SDs and are more likely to have recently duplicated gene families with higher copy number when compared to non-African samples. A comparison to a resource of 563 million full-length Iso-Seq reads identifies 201 novel, potentially protein-coding genes corresponding to these copy number polymorphic SDs.

## INTRODUCTION

The first draft sequences of the human genome ^1^ revealed a surprising degree of high-identity duplications dispersed both interchromosomally and intrachromosomally. Segmental duplications (SDs) have been operationally defined as blocks of homologous DNA greater than 1 kbp in length with >90% sequence identity ^2^. In humans, ∼60% of the pairwise alignments are interspersed and separated by more than 1 Mbp within a given chromosome or mapping to the pericentromeric or subtelomeric regions of nonhomologous chromosomes ^3,4^. Because of their size and high degree of sequence identity, SDs have been some of the last regions of the human genome to be fully resolved ^5^. Originally estimated at 5% of the genome, the relative proportion within the telomere-to-telomere (T2T) genome has increased to ∼7%, especially as the acrocentric regions of the short arms of human chromosomes have become fully characterized at the sequence level ^6,7^.

SDs contribute disproportionately to human genetic diversity because of their potential to drive unequal crossing over (aka non-allelic homologous recombination or NAHR). Based on sequence read-depth analysis of the short-read sequencing data from the 1000 Genomes Project (1KG) ^8^, for example, we estimated that 50% of all copy number polymorphisms in the human species >1 kbp in length map to SDs—an ∼10-fold enrichment ^9^. Importantly, almost all copy number polymorphic genes in the human species map to these particular regions of the genome ^10^. Such copy number polymorphic genes have been strongly implicated in a variety of human diseases ranging from immune/autoimmune (*FCGR*) ^11–13^, neurological (*C3/C4*) ^14,15^, to coronary heart disease (*LPA*) ^16,17^. More recently, it has become apparent that genes embedded within SDs play an important role in the evolution of our species, including the expansion of the human frontal cortex (*SRGAP2C* ^18^, *ARHGAP11B* ^19^, *TBC1D3* ^20^), adaptation to starch-rich diets (amylase ^21,22^), or even the development of color vision within the primate lineage (green and red opsins ^23^).

Notwithstanding their importance, understanding the genetic diversity of these more complex regions of the genome has been challenging. Most efforts have focused on estimating copy number by mapping short-read data back to a singular reference to discover copy number variant (CNV) regions ^24–26^. Such investigations are problematic for two reasons. First, they tell us almost nothing about the location or the structure of the CNVs or the actual protein-coding potential of the underlying genes. Second, they introduce a reference bias since, until recently, the human reference genome was incomplete—with gaps enriched precisely over the most CNV regions. Advances in long-read sequencing technology over the last four years, however, have made these regions accessible for the first time ^6,27,28^. In particular, the development of PacBio HiFi (high-fidelity) sequencing technology and associated assembly algorithms ^29,30^ has meant that most SD regions can be fully sequence resolved at the haplotype level. In this study, we sought to investigate the population genetic diversity of SDs by focusing on 170 human genome assemblies for which HiFi sequence data had been collected as part of the Human Pangenome Reference Consortium (HPRC) and Human Genome Structural Variation Consortium (HGSVC) ^10,31^.

## RESULTS

### Distribution of shared versus polymorphic SDs

In this analysis, we initially considered a diverse set of 106 human samples (212 haplotype assemblies), all of which originated from the 1KG and for which sufficient HiFi sequence data had been generated as part of previous efforts ^10,31^. This included 47 HPRC (all trio binning assemblies using parental short reads) and 53 HGSVC (14 trio and 39 non-trio) samples. We sequenced and assembled all genomes using the same assembly algorithm, hifiasm (v0.14/v0.16), which had been shown previously to accurately resolve most (although not all) SD regions ^10,32^. Because of the potential for assembly collapse, we restricted our analysis to 1KG samples where matched Illumina short-read sequence data were available, and the genomes passed QC and were all assembled with the same algorithm (Methods). SDs are particularly prone to assembly errors or collapses and this procedure both harmonized the results and allowed for all duplicated sequence to be validated by Illumina read-depth analysis. In total, we analyzed 170 independent genome assemblies and identified SDs (>1 kbp and >90%) from 85 human specimens representing 38 African and 47 non-African samples (Supplementary table 1).

To investigate how SD patterns vary among human genomes, we mapped SDs back to the T2T human reference genome (T2T-CHM13) classifying events as either known or new with respect to that reference and then assessed whether they were shared or unique among the 170 human haplotypes. We used these data to estimate the allele frequency of inter and intrachromosomal duplications creating a pangenome representation of human SDs (Figure 1). Because of the difficulties in both assembly and mapping of acrocentric SDs, we excluded all short arms or acrocentric chromosomes from this analysis. In total, we identified of 2,742 intrachromosomal and 4,772 interchromosomal nonoverlapping SD regions, constituting 6.1% of the genome or 173.21 Mbp (150.12 Mbp and 73.95 Mbp for intra and interchromosomal SDs, respectively; 50.86 Mbp overlapped between intra and interchromosomal SDs) based on the genomic coordinates of the T2T-CHM13 genome.

**Fig. 1.**
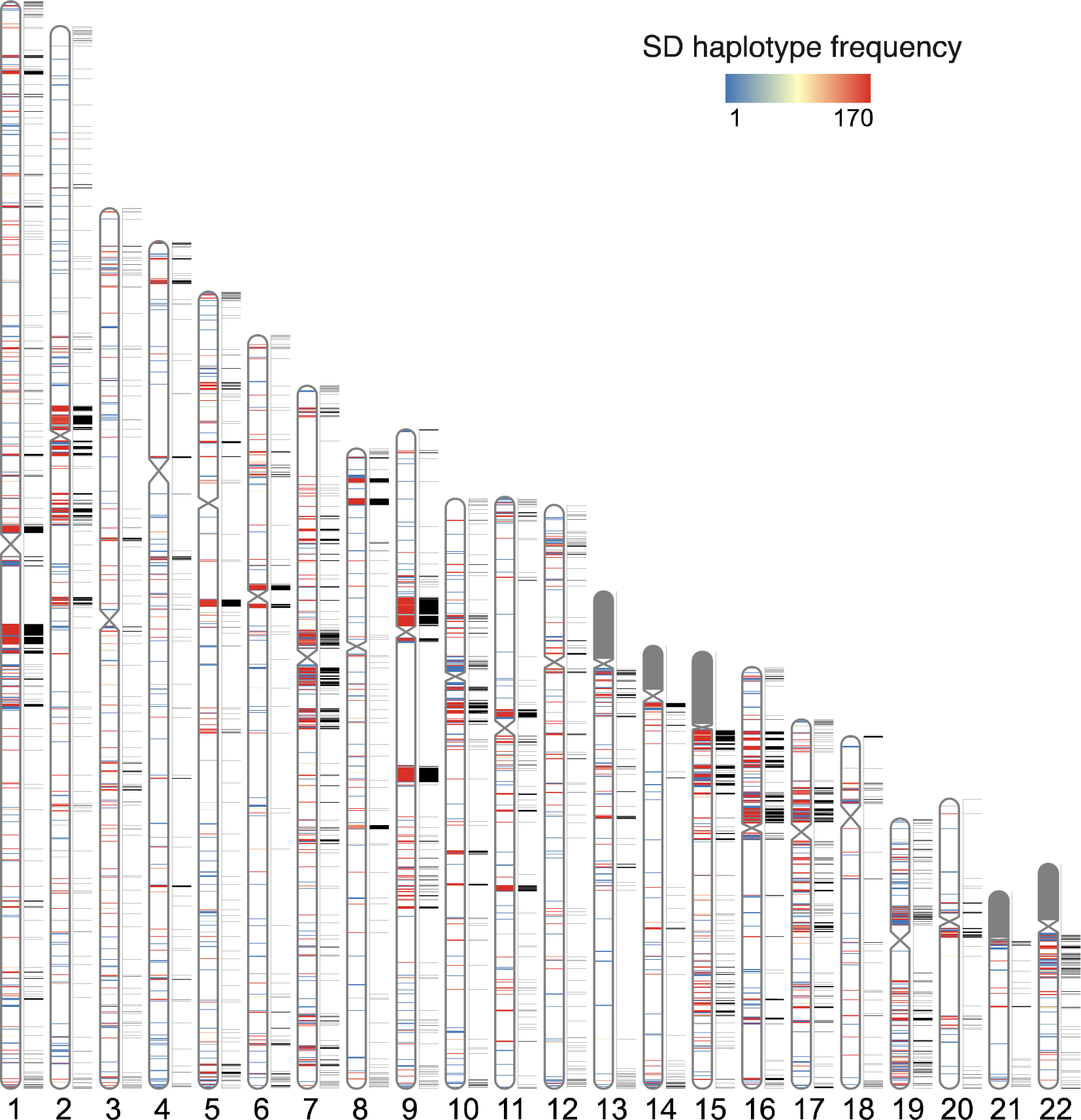
Pangenome representation of human segmental duplications (SDs) Haplotype frequency distribution of intrachromosomal SD content from HPRC and HGSVC haplotype genome assemblies (n=170). SDs are colored by the haplotype frequency. SD content on the p-arms of acrocentric chromosomes (chr13, chr14, chr15, chr21, and chr22) was excluded due to assembly errors and potential chromosomal misassignment compared to other autosomal chromosomes. The known SDs of T2T-CHM13 are shown in black next to the ideograms on each chromosome.

Compared to the T2T-CHM13 human genome, we classify 47.4 Mbp overall as newly discovered with an estimated rate of accumulation of 408.3 kbp SDs being added for each additional sequenced and assembled human genome (Figure 2). The majority of these novel SDs map intrachromosomally (41.7 Mbp) although we classify 7.4 Mbp as interchromosomal—mapping primarily to subtelomeric and pericentromeric regions of the human genome (24.6%, p-value < 0.01, odds ratio = 2.39). Overall, a greater fraction of interchromosomal SDs (78.4%) is fixed when compared to intrachromosomal events (59.7%). With respect to intrachromosomal duplications, we find the majority of novel SDs tend to occur in close proximity to previously known SDs (empirical p-value < 0.01, permutation test) although we do note that certain chromosome arms and regions (e.g., p-arms of chromosomes 8, 10, 16, 17, 19 and q-arms of chromosomes 1, 15, 22) show an excess of these rarer SDs (Figure 1 and Supplementary table 2). As expected from other structural variation studies, the accumulation of novel SDs shows an asymptotic relationship with increasing sample size and the accumulation is greater for the additional African samples owing to their overall increased genetic diversity (Figure 2).

**Fig. 2.**
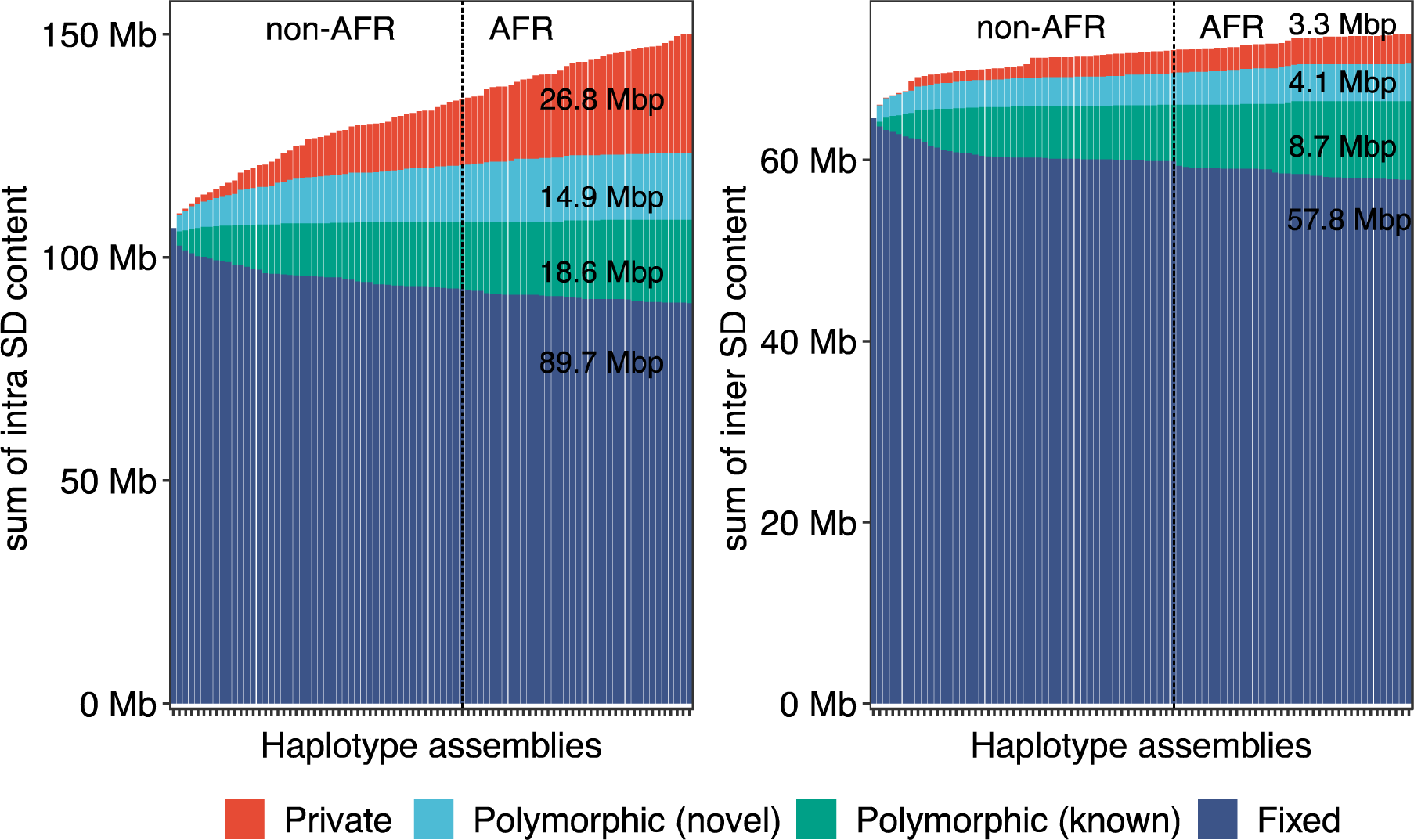
Cumulative sum of SDs by frequency. Bar plot displays the cumulative sum of SD content by adding genomes (from left to right) for intrachromosomal and interchromosomal SDs. Four SD frequency categories are considered: “Fixed” are SDs present in all 170 human genome assemblies (i.e., conserved in all samples); “Polymorphic (known)” are SDs in the reference genome (T2T-CHM13) that are not fixed; “Polymorphic (novel)” refers to SDs observed in two or more HPRC/HGSVC assemblies yet not present in T2T-CHM13; “Private” is an SD found in one sample. Samples are grouped by non-African (non-AFR) and then African (AFR) genetic ancestry due to the expected increased diversity among the latter.

### Sequence properties of polymorphic and rare SDs

Among the polymorphic SDs, we further distinguished two groups: rare SDs observed in up to five human genomes (<3% allele frequency) and common SDs observed between 6-20 times (∼3-10% allele frequency). We find that rare SDs tend to be longer and have higher sequence identity between SD pairs when compared to common SDs (empirical p-value < 0.01, permutation test) (Figure 3A and Supplementary figure 1). These features are consistent with a more recent origin for the majority of rare SDs; however, we note that highly divergent SDs are still identified that occur at a low frequency in the human population (Figure 3A). Notably, we find that low-frequency SDs also tend to be more distant from known SDs than those with a higher allele frequency in the population suggesting that the interspersion process characteristic of ape genomes is still ongoing in the human population.

**Fig. 3.**
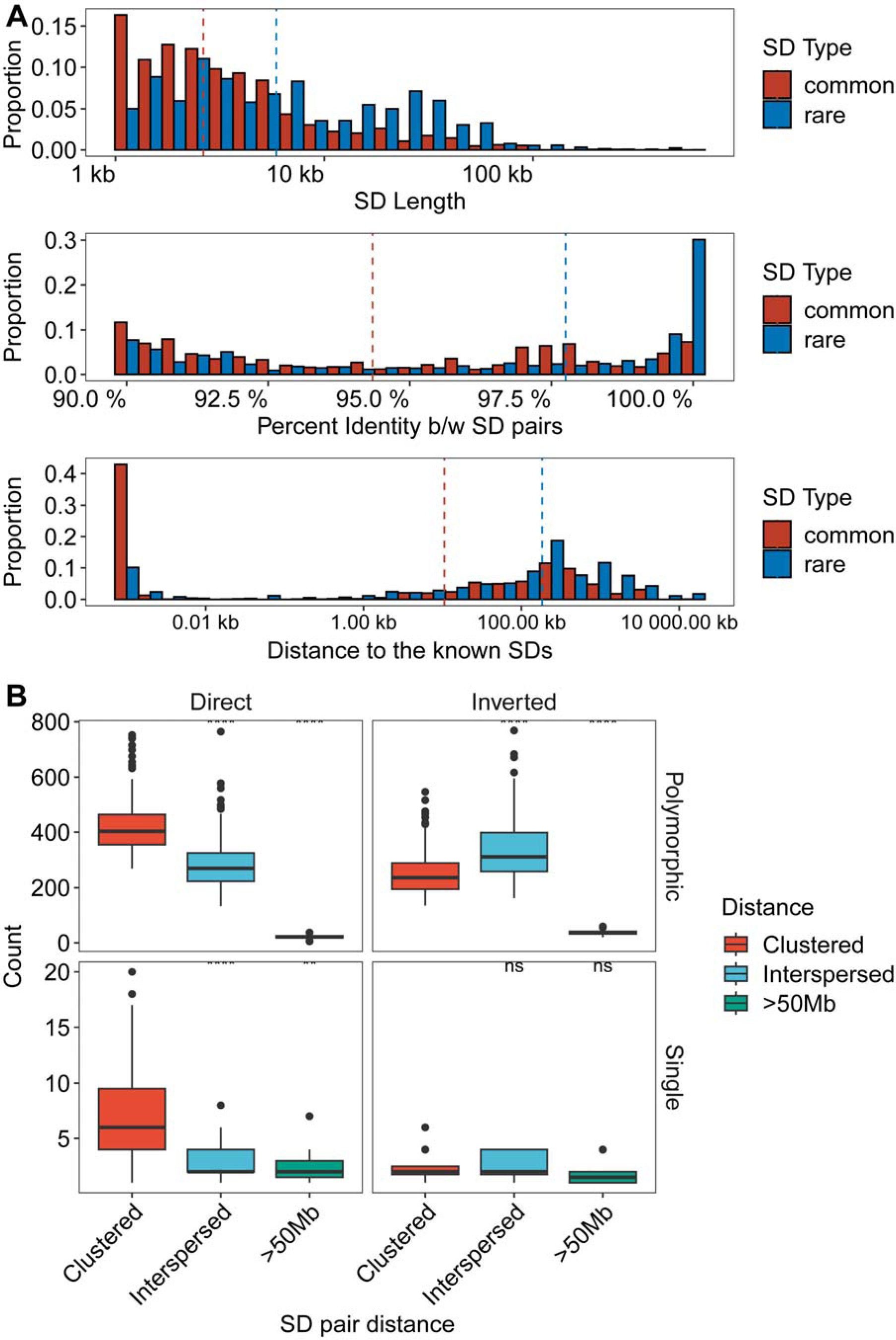
Sequence properties of polymorphic versus rare SDs. **(A)** Histogram comparing the sequence identity and length of rare and common SDs (see Supplementary figure 1 for polymorphic SDs with more subclassified haplotype frequencies). **(B)** Orientation and pairwise dispersion of polymorphic and singleton SDs. Each data point represents haplotype assembly, and their counts of clustered, interspersed (>1 Mbp apart), and distant (>50 Mbp apart) SDs. Left and right panels summarize the SDs in direct or inverted orientation while the top and bottom panels contrast polymorphic vs. singleton SDs.

We also considered the orientation and configuration of singleton (single occurrence in a genome validated by read depth) versus polymorphic (not fixed in all human genome assemblies with allele frequency below 90%) SDs. We find that the vast majority (89.1%) of all SD singletons are clustered irrespective of whether they exist in a direct or an inverted orientation (Figure 3B; Supplementary table 3). We find that the proportion of polymorphic SDs classified as interspersed (SD pairs separated by more than 1 Mbp) increases in approximately equal proportions between inverted and directly orientated SDs (Figure 3B). However, interspersed polymorphic SDs favor an inverted orientation (p = 9.6 x 10^-10^, odds ratio = 1.99, Fisher’s exact test). Figure 4 depicts examples of the structure of rare SDs in an inverted orientation.

**Fig. 4.**
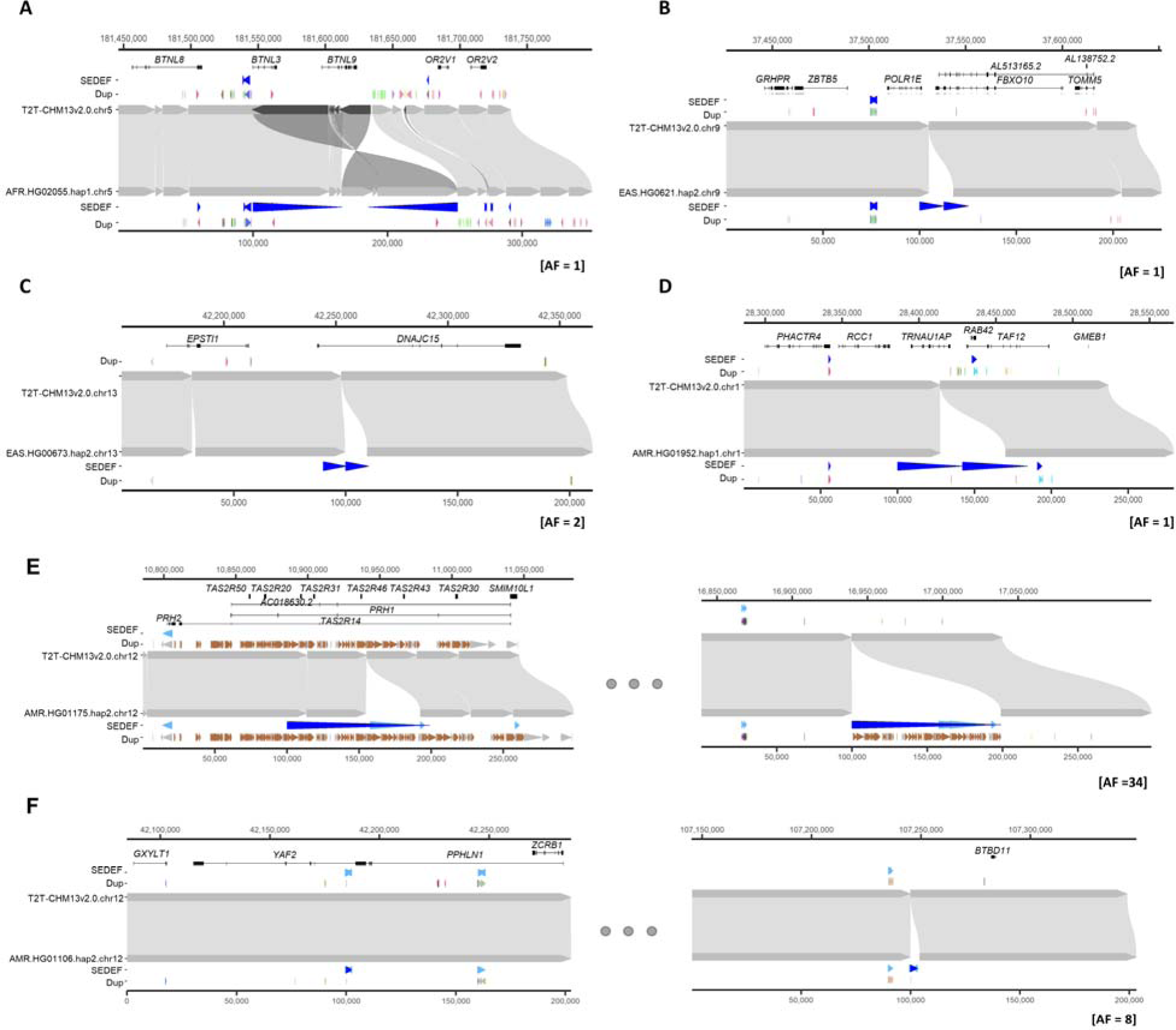
Examples of clustered (A-D) and interspersed (E-F; >1 Mbp apart) SDs associated with genes. In each plot, the top represents the T2T-CHM13 genome aligned to bottom, new genome assemblies. **(A)** Clustered duplication with inverted orientation (65.8 kbp; with allele frequency [AF] = 1) found in chr5 and **(B-D)** clustered and tandem duplications (12.6, 10.3 and 42.3 kbp; with AF of 1, 2 and 1, respectively) in chr9, chr13 and chr1. **(E-F)** Interspersed duplications of chr 12 (98.9 and 2.5 kbp; with AF = 34 and 8) showing duplicated regions in left and right panels. The gene track of the T2T-CHM13 genome assembly is shown at the top, followed by SDs predicted by SEDEF and the respective direction indicated by blue arrowheads. The DupMasker track shows the duplicon structure.

### Gene content and population differences copy number

Based on the current gene annotation of the T2T-CHM13 genome assembly, we estimate that there are 1,156 duplicated protein-coding genes (standard deviation = 49) per diploid. Considering all SDs identified in the 170 human genomes, we estimate that 1,340 protein-coding genes are duplicated to copy number four in at least one sample. Of note, 173 of these correspond to single-copy genes in the T2T-CHM13 reference (Methods; Supplementary table 4). The majority of the low-frequency SDs are incomplete with respect to their ancestral gene model, often involving a subset of the original exons. We caution, however, that incomplete SDs do not guarantee that the duplicates are pseudogenes ^18,19,33^.

Next, we considered multicopy SDs based on gene content, grouping 1,095 multicopy genes in the T2T-CHM13 reference into 314 gene families. As a control for potential assembly artifacts, we orthogonally evaluated gene copy by correlating (R-squared = 0.94) assembly gene copy by predicted copy number from Illumina short-read sequencing read depth (Supplementary figure 2). Using the dispersion index as a metric, we identified the 25 most variable gene families in the human genome and contrasted them with the 25 least variable (Figure 5A-B and Supplementary figure 3). As expected, higher copy number gene families (10-50 members) were among the most variable in the human population while the most invariant typically were fixed at four or six copies (diploid copy number). The least-variable gene family, *HYDIN*, includes the human-specific duplication *HYDIN2*, which gained neural expression by adopting a new promoter ^34^. Similarly, the *RGPD3* family is the eighth least variable based on its copy number and includes two human-specific copies that we find to be under selection ^35^. A gene ontology analysis showed that highly variable gene copy associated with female pregnancy (*PSG*), amylase activity (*AMY2*, *AMY1*) and immune response (defensin, *KIR2DL*) and unknown biological function while fixed copy number SD genes were particularly common among Kruppel-associated box (KRAB) zinc-finger proteins (KRAB-ZFPs) and genes associated with metabolic process (*CYP1*, *CYP2* and *CYP4*) (Supplementary figure 4). However, even among high copy number genes, both variable and invariant members are observed.

**Fig. 5.**
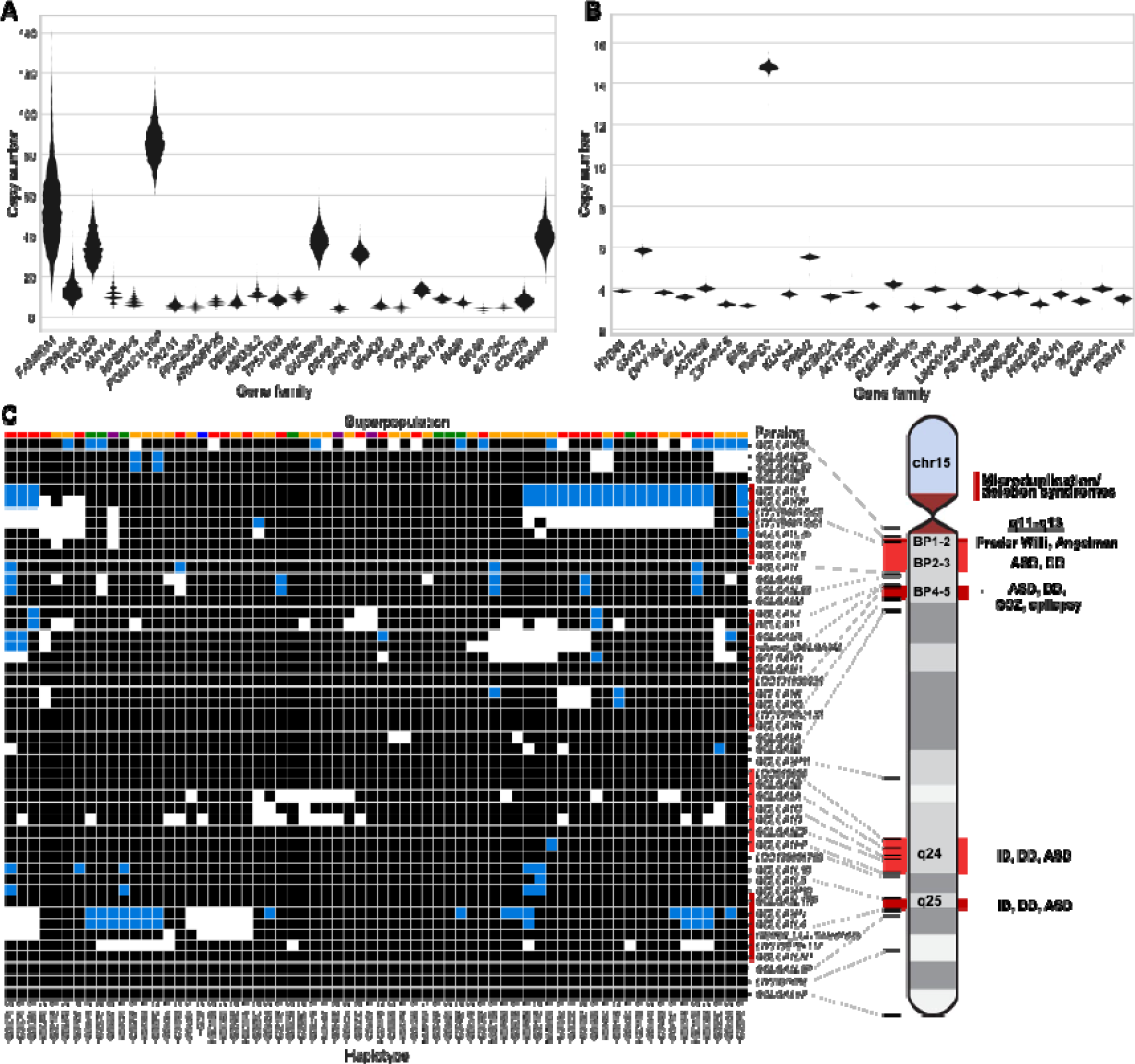
Variable copy number of duplicated genes. **(A-B)** Gene families with highly variable (A) and nearly fixed (B) copy number are displayed. Gene families are selected and ordered by dispersion index, requiring an average diploid copy number greater than three. **(C)** Estimated copy number of *GOLGA6/8* paralogs in each assembled haplotype, based on assembly alignments (white:0, black:1, blue:2). The continental population groups for each haplotype are indicated by color above each column (Africa: gold, East Asia: green, South Asia: purple, Europe: blue, the Americas: red). ASD: autism spectrum disorder, DD: developmental delay, ID: intellectual disability, SCZ: schizophrenia.

Using these highly contiguous assemblies, we are now able to assign copy number polymorphism to specific paralogs in addition to assaying copy number on the level of gene families. For example, for one of the most variable gene families in the human genome (*GOLGA6/8*), we analyzed the copy number variation of each *GOLGA* paralog across our assembled haplotypes (Figure 5C). To avoid false duplicates, we only included haplotypes with no assembly breaks within 30 kbp of the *GOLGA* paralog. Of the named protein-coding paralogs, only *GOLGA6L2*, *GOLGA8M*, *GOLGA8H*, *GOLGA8N*, and *GOLGA6B* are fixed at a single copy across all haplotypes, while four are variable but never deleted, and the remaining 18 are deleted or absent in some haplotypes. This paralog specificity identifies those five single-copy genes as higher-priority candidates for functional analysis, given they are fixed in the human population. *GOLGA* paralogs mediate pathogenic microduplications and deletions at 15q11-13, q24, and q25 causing forms of intellectual delay, including Prader-Willi syndrome ^36–40^. While we observe multi-gene deletions in these regions (Figure 5C), including genes such as the human-specific fusion gene *CHRFAM7A* whose deletion has been implicated in Alzheimer’s disease pathology ^41^, none of these deletions extend beyond the SD into the unique critical regions for named syndromes in our samples.

During this gene analysis, we noticed that samples of African ancestry tended to show overall higher copy number for multicopy SDs. We tested this more formally in three ways. First, we compared the intra and interchromosomal content irrespective of gene content between genomes of African or non-African origin. Genomes of African origin harbor significantly more intrachromosomal SDs (p=1.6 x 10^-6^, Mann-Whitney U test) (Figure 6A). Next, we examined the copy number of gene families by two methods: by counting assembled paralogs in our long-read assemblies and by using read depth to estimate copy number in a larger cohort with short reads (n=2,196). In the long-read assemblies, we tested the 90 protein-coding gene families with variable copy number (dispersion index ≥ 0.1) and mean copy number greater than two for African or non-African samples. Seventeen gene families showed shifted copy number distribution, and 16/17 showed the same effect by read-depth analysis (Mann-Whitney U test, Benjamini-Hochberg corrected p ≤ 0.05). Consistent with the increase in intrachromosomal SDs, for 13/16 gene families (81%), the copy number distribution is higher in African than non-African samples, as shown in Figure 6B (binomial test, p = 0.01). Finally with the larger sample of high-coverage Illumina data from unrelated individuals in the 1KG, excluding highly admixed populations (n=2,196), we considered the 1,171 gene families with dispersion index ≥ 0.1 and mean copy number greater than two in African or non-African samples (Supplementary figure 5). Population-differentiated copy numbers are observed in 263 gene families (Mann-Whitney U test, Benjamini-Hochberg-corrected p < 0.05), with 164/263 (62%) shifted towards higher copy number in African samples (p = 0.00004, binomial test). The gene families with largest shifts (greater than 15%) are shown in Figure 6C, with 17/22 (77%) shifted towards higher copy number in African samples. All statistically significant gene families are shown in Supplementary figure 6. From the assembly test of copy number differentiation, only *GUSBP3* did not replicate in the larger read-depth cohort.

**Fig. 6.**
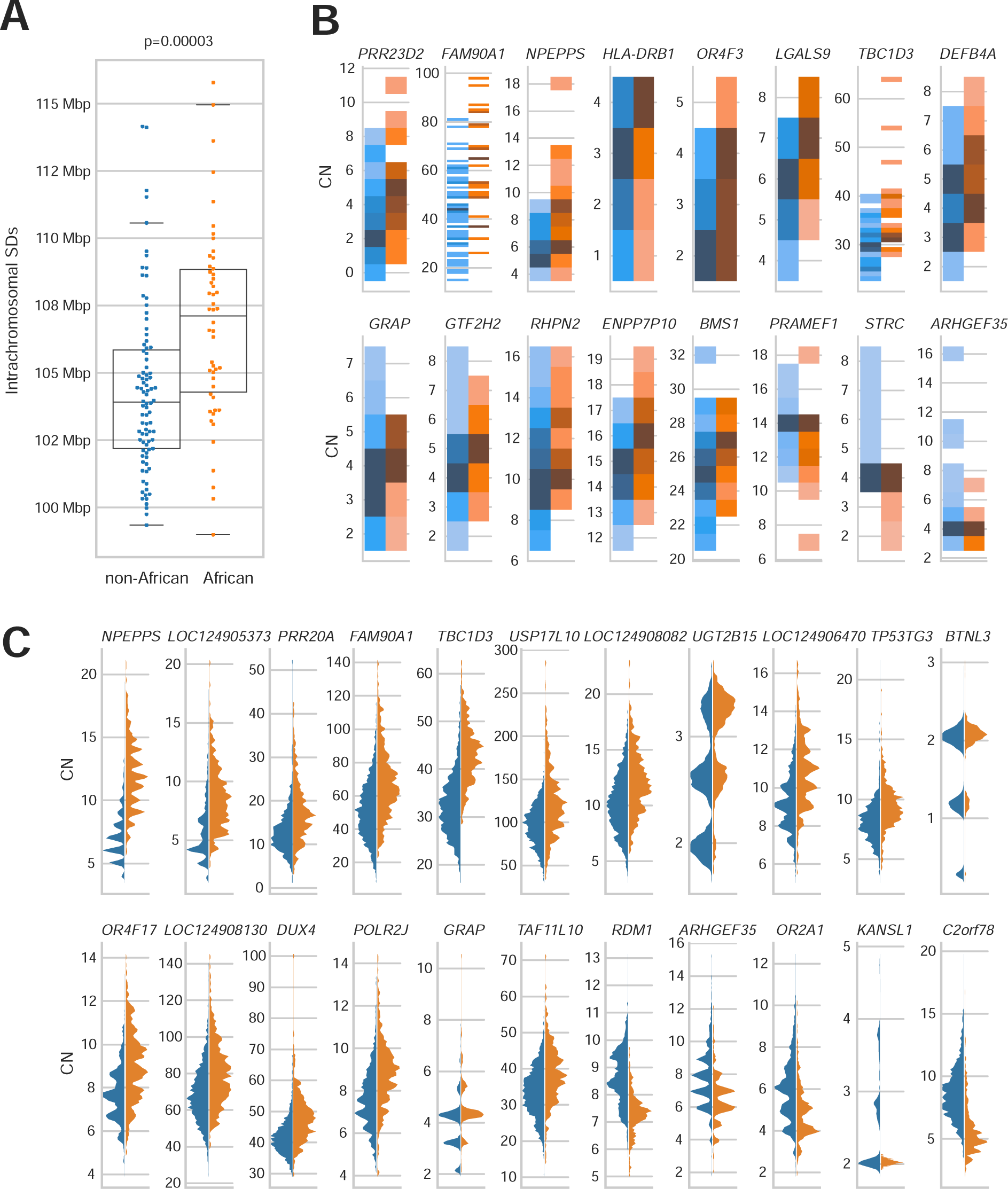
African vs. non-African SD copy number variation. **(A)** Proportion of intrachromosomal SD content between African and non-African populations. African genomes have a higher SD content compared to non-African genomes, and the difference is significant for intrachromosomal SDs. **(B)** Gene family copy number variation between populations. Gene families with significant copy number differences between African and non-African populations are shown (Mann-Whitney U test, Benjamini-Hochberg adjusted p-value <0.05), excluding *GUSPB3*, which did not replicate in the larger cohort. Gene copy number (CN) was estimated from the assemblies by whole-genome alignment; 13/16 gene families average higher copy number in individuals of African ancestry (binomial, p = 0.01). **(C)** Gene copy number evaluated by Illumina read depth. The 22 gene families with the largest distribution shift are shown.

### Genic potential of polymorphic SDs

We sought to assess the transcriptional potential of the structurally polymorphic SDs identified in this study. Because of the high degree of sequence identity among the SDs, gene annotation has been difficult with standard RNA-seq datasets because short reads map equally well to distinct loci. This is especially true for copy number polymorphic genes where individual copies are >99% identical and can range in copy from 5-40 among different individuals in the population ^42^. To address this limitation, we assembled a long-read Iso-Seq resource of 563 million full-length non-chimeric (FLNC) cDNA sequences generated from 241 libraries and 67 distinct tissues (Supplementary table 5). We mapped each FLNC read both to the T2T-CHM13 genome and a pangenome of 170 human genomes searching specifically for FLNC reads that mapped better to the pangenome. Specifically, we required at least 99.9% sequence identity to an assembled haplotype and less than 99.7% gap-compressed identity to T2T-CHM13—below the expected allelic divergence for most protein-coding regions of the genome. We focused on putative protein-coding genes and constructed 7,081 gene models for cDNA alignments spanning 476 Mbp of T2T-CHM13.

We used these reference-divergent cDNA reads that matched better to other assembled haplotypes (n=1,279,037) to predict protein-coding genes (Figure 7A). Each additional human haplotype contained an average of 46 protein-coding gene predictions that showed more than 1% divergence from T2T-CHM13 reference annotations (range 13– 77), highlighting the importance of additional human genome references to fully assess human genic variation. To count novel gene annotations across haplotypes, we grouped genes/transcripts into gene families by counting only predictions from the haplotype with greatest number of novel paralogs. This resulted in a total count of 260 putative novel protein-coding genes from 206 gene families. Of these 260 genes, 183 mapped to SD regions, 18 genes mapped to SD regions for at least one sample but not the T2T-CHM13 reference, and the remaining 59 genes mapped to unique sequence (not SDs). Gene ontology biological process enrichment analysis of these genes compared to the background of protein-coding genes within the 476 Mbp of sequence examined yielded 13 significantly enriched driver terms, largely related to immunity: positive regulation of leukocyte mediated immunity, antigen processing and presentation of endogenous peptide antigen, rRNA metabolic process, symbiont entry into host, T cell extravasation, regulation of deoxyribonuclease activity, leukocyte cell-cell adhesion, regulation of type II interferon production, regulation of lymphocyte activation, dendritic cell differentiation, detection of bacterium, peptide antigen assembly with MHC class II protein complex, and positive regulation of cell-cell adhesion (Benjamini-Hochberg adjusted p < 0.05). Twenty of the novel genes belong to the immunoglobulin superfamily ^43^ while ten are within core duplicons, a group of loci hypothesized to drive the evolution of interspersed larger SD blocks ^44^ during ape evolution. Only 17.5% (28/160) of these predicted protein sequences had previously been submitted to GenBank.

**Fig. 7.**
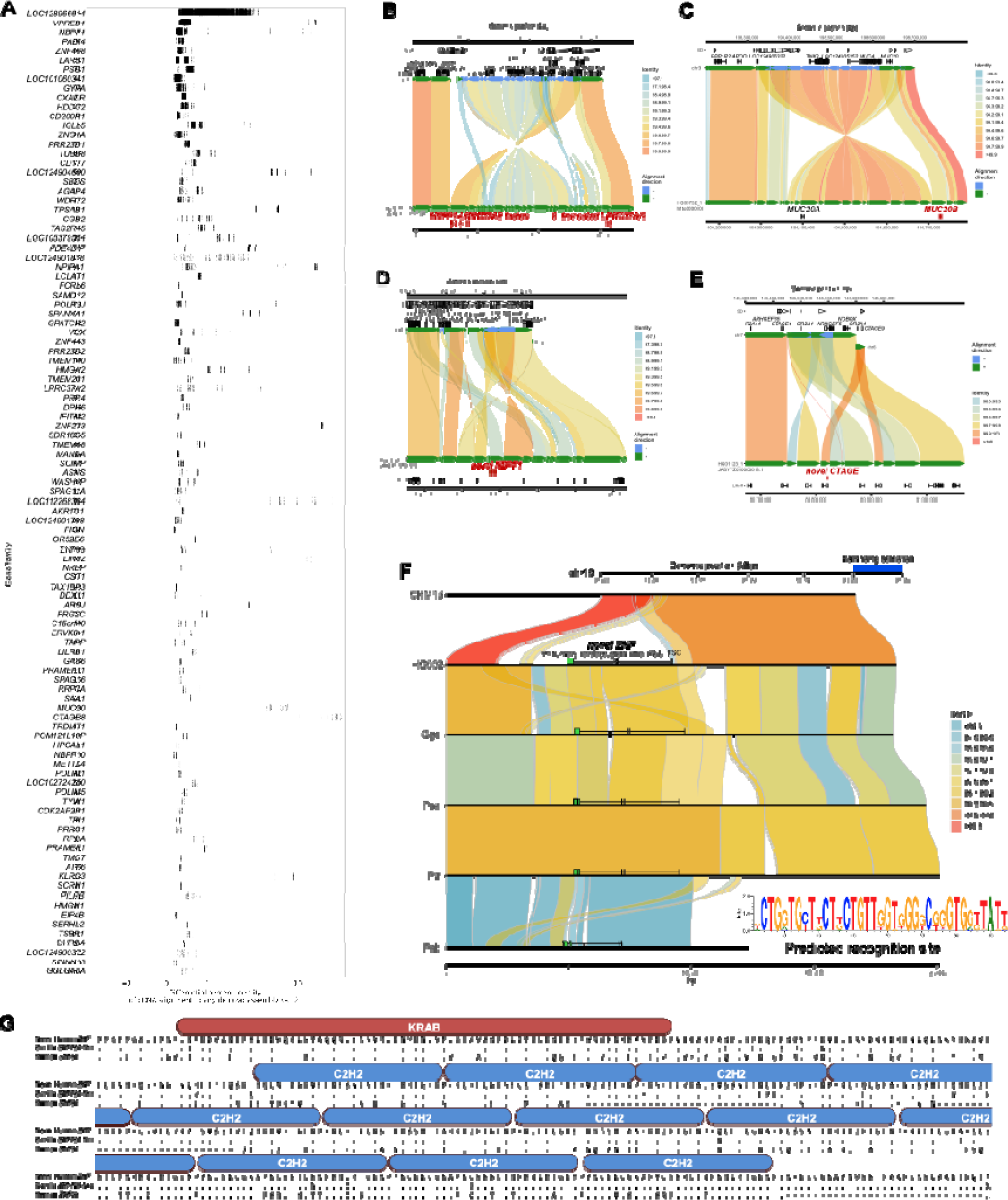
Discovery of novel gene/transcripts in rare and polymorphic SD regions. **(A)** Examples of copy number polymorphic gene families where FLNC generated from Iso-Seq map better to the pangenome than to the T2T-CHM13 human genome reference. **(B-E)** Selected haplotypes containing novel gene predictions for *MUC20, NBPF1, CTAGE*, and *LRRC37A* compared to T2T-CHM13 reference where there is FLNC transcript support. Alignment color indicates percent identity. **(F)** Comparison of T2T-CHM13 (top) and HG002 maternal haplotype (bottom) depicts 48 kbp polymorphic SD region present in 66/170 haplotypes. Nonhuman apes all carry a copy of the duplicated sequence. *ZNF* predicted recognition site shown (inset). **(G)** Comparison of the novel *ZNF* to its best human match (*ZNF98*, 68% identity), and the most similar existing primate annotation (low-quality protein *ZNF724*-like in gorilla, 95% identity). ProSite-predicted KRAB-ZNP is shown above the sequence.

Notable examples of novel gene annotations include additional copies of *MUC20*, *GSTM*, *TUBB8*, *SIRPB1*, *GOLGA8*, *LRRC37A*, *NBPF1*, *CTAGE*, and *UPK3BL1* (Figure 7B-E) ^45–47^. The paralogs with the lowest identity cDNA sequence compared to T2T-CHM13 often have modified isoform structures, predominately modified N- or C-termini due to the structural rearrangements that led to their formation. For example, the 17q21.31 *KANSL1* inversion haplotype H2 includes a partial *KANSL1* duplication, which acts as an alternate 5’ promoter for *LRRC37A/2*, producing a putatively protein-coding transcript with 39 novel amino acids at its N-terminus followed by sites 870 to 1700 of the canonical *LRRC37A/2* isoform (Figure 7B). The same haplotype also encodes a *NSFP1-LRRC37A2* fusion transcript, which maintains an open reading frame, predicted to produce a protein with the first 492 amino acids of *NSF* followed by amino acids 41-903 belonging to the core duplicon gene *LRRC37A2* (Supplementary figure 7).

We discovered a novel expressed copy of *MUC20* in the paternal haplotype of HG03732 within a 214 kbp inversion at chr3q29 that duplicates 73 and 37 kbp on its edges (Figure 7C); its expression is supported by full-length cDNA from 14 Iso-Seq libraries from chondrocytes, soft tissue, left colon, induced pluripotent stem cells (iPSCs), human embryonic stem cells, and fibroblast cell lines. Structural diversity at 3q29 has been documented in prior work ^43^ but the expression and gene model of the additional *MUC20* paralog has not yet been reported to our knowledge. Two assembled haplotypes have a complex rearrangement at chr1p36.13 that creates 86 kbp of additional sequence compared to T2T-CHM13, including an additional copy of *NBPF1* supported by two Iso-Seq reads from brain tissue and a mammary epithelial cell line (Figure 7D). In the paternal haplotype of HG01123, an additional copy of *CTAGE* is created by a 16 kbp insertion from chr6q23.2 into chr7p35 in the context of a 59 kbp duplicated inversion, with expression detected with a single read from each of four Iso-Seq libraries from promyeloblast cells and iPSCs (Figure 7E). Even among *HLA* genes, whose polymorphisms have been extensively documented due to their clinical significance, we identified 62 novel alleles across seven distinct genes not currently represented in GenBank or the IPD-IMGT/HLA database (*HLA-A, -B, -C, -G, -DQB1, - DRB1, -DRB5*).

During this analysis, we identified low-identity alignments to *ZNF724* corresponding to a novel KRAB-ZNP absent from the T2T-CHM13 reference. This duplicate gene, provisionally named *ZNF972*, has only 69% identity to its best-matching annotated human gene, *ZNF98* (Figure 7). *ZNF972* cDNA reads were found in 18 (6%) Iso-Seq libraries and correspond to a 48 kbp region within the chr19p12 ZNF cluster, not present in previous human reference genome assemblies (GRCh38 and T2T-CHM13), though it exists in 35.9% of the assembled human haplotypes we analyzed. This region is also present in *Pan*, *Gorilla*, and *Pongo* genomes, but an orthologous gene has only been annotated in *Gorilla* as *ZNF972* coding sequence. Its open reading frame is disrupted in Pan and Pongo relative to Gorilla and human (Figure 7F-G). Thus, ZNF972 is an example of an ancestral ape duplicated gene that is still present in Gorilla and is present in a subset of humans but likely pseudogenized in other ape lineages.

## DISCUSSION

The last two decades of human genomics research have shown that SDs play an important role in human health and evolution, contributing to genetic diversity, adaptation, selection, genomic instability, and susceptibility to disease ^18,19,33,48–53^. Despite their importance, understanding how humans vary with respect to this structural feature of our genomes and its potential functional consequence has always been challenging in large part because the size and high sequence identity of SD repeats have made interrogation of these regions and ∼1000 protein-coding genes mapping within almost impossible with traditional sequencing and genotyping approaches. As a result, most genome-wide association studies as well as genome-wide surveys of selection, gene regulation (ENCODE), and transcription (GTEx) have explicitly excluded the most identical SDs from study ^54,55^. Even early long-read sequencing-based approaches failed to adequately resolve these particular regions ^31,53^. The advent of PacBio HiFi sequencing data ^56^ along with improved assembly algorithms has fundamentally changed the calculus ^29,30,30,57^. The sequence accuracy of HiFi data (>99.9%) meant that paralogs and alleles could be fully resolved in a phased genome assembly making these regions systematically accessible for the first time ^10,57,58^. In this study, we took advantage of HiFi data generated as part of the HGSVC and HPRC to analyze SDs in a total of 170 genome assemblies and compare the results to a complete human reference genome (T2T-CHM13). We harmonized the data using the same assembly algorithm and validated copy number in individual genomes using Illumina whole-genome sequence data to reveal the location and structure of copy number of SD variation at a population level for the first time. Thus, this pangenome representation provides one of the first glimpses of human structural diversity of SDs genome-wide.

While SDs have been known to be enriched in copy number polymorphisms ^25,59,60^, the phased genome assemblies allow us to quantify, map, and compare this variation revealing some unexpected findings. Our analysis of 170 genomes identifies 76.4 Mbp of variable versus 147.5 Mbp of invariant SD DNA—the latter may be more likely to harbor genes that will be functionally constrained ^18^. Although fundamentally different in nature, the amount of genetic variation in this 6% of the genome in these 85 individuals is comparable to the estimated 84.7 million single-nucleotide polymorphisms discovered genome-wide from sequencing the 2,500 individuals from the 1KG ^61^. We find that intrachromosomal SDs are twice as likely to be polymorphic when compared to interchromosomal SDs, although we should caution that we excluded the acrocentric regions of human short arms from this analysis where we anticipate rampant ectopic recombination and interchromosomal copy number variation to occur ^62^. While most of the 41.4 Mbp of novel SDs (with respect to the finished T2T-CHM13 genome) occur in close proximity to existing regions of SD, we discovered novel sites of interspersed duplications. Remarkably, such interspersed rare SDs are more likely to be configured in an inverted orientation minimizing predisposition to large-scale microdeletions although potentially promoting rare inversion polymorphisms in the population ^32^. Also, we note that certain chromosome arms, including chromosome 1q, 8p, 10p, 15q, 16p, 17p, 19p, and 22q, appear enriched for novel SDs—the basis for this chromosomal bias is unknown.

From a population genetics perspective, it is noteworthy that samples of African ancestry show significantly greater intrachromosomal SD content when compared to 1KG populations belonging to other continental groups. This translates into an overall higher gene copy number for duplicated genes (Figure 6)—an observation we confirmed both by genome assembly as well as Illumina read-depth analyses (Methods). While increased variance in copy number would be consistent with the overall 15-20% increase in genetic diversity and greater population substructure that has been reported for populations of African ancestry ^63^, an overall higher copy number for duplicated gene families, especially those related to environmental interaction (e.g., drug detoxification, immunity), may have provided ancestral human populations with greater buffering capacity and adaptive potential but also greater susceptibility to NAHR-mediated rearrangements. It is interesting that read-depth sequencing analysis of some archaic hominins such as Denisova have suggested overall higher copy number for many gene families when compared to modern humans ^64^.

There are several limitations of the current study. First, we sampled only 85 individuals (170 human genomes) and this represents only a small swath of potential human genetic diversity. As more human genomes are sequenced and pushed toward T2T status ^65^, a more complete picture of human genetic diversity will begin to emerge. This will include population-specific paralogs and insights into the mechanisms underlying the formation of interspersed SDs as well as the role of SDs in driving ectopic recombination of acrocentric short arms in the human population ^66^. Similarly, our attempt to identify novel genes using a deep resource of human Iso-Seq data should be regarded only as a starting point. The challenge especially for assessing rarer SDs that harbor duplicated genes is that full-length cDNA was derived from different individuals from those whose genomes were sequenced and assembled ^10,67^. Genomic resources, such as those being generated from the SMaHT (Somatic Mosaicism across Human Tissues) initiative where donor-specific assemblies and Iso-Seq data from different human tissues from the same source are gathered, will be required ^68,69^. Such matched transcription and assembly data from the same donor will provide a clearer picture of the transcription as well as the tissue specificity of the thousand genes mapping to human duplicated sequence. Ultimately, functional characterization will be required to confirm the missing protein-coding copy number polymorphic genes in our genome.

## METHODS

### PacBio HiFi sequence production

#### University of Washington

Isolated DNA was sheared using the Megaruptor 3 instrument (Diagenode) twice using settings 31 and 32 to achieve a peak size of ∼15–20Lkbp. The sheared material was processed for SMRTbell library preparation using the Express Template Prep Kit v2 and SMRTbell Cleanup Kit v2 (PacBio). After checking for size and quantity, the libraries were size-selected on the Pippin HT instrument (Sage Science) using the protocol ‘0.75% agarose, 15–20 kb high pass’ and a cut-off of 14– 15Lkbp. Size-selected libraries were checked by fluorometric quantitation (Qubit) and pulse-field sizing (FEMTO Pulse). All cells were sequenced on the Sequel II instrument (PacBio) with 30 h video times using version 2.0 sequencing chemistry and 2 h pre-extension. HiFi/CCS analysis was performed using SMRT Link (v.10.1) using an estimated read-quality value of 0.99.

#### The Jackson Laboratory

High-molecular-mass DNA was extracted from 30Lmillion frozen pelleted cells using the Gentra Puregene extraction kit (Qiagen). Purified gDNA was assessed using fluorometric (Qubit, Thermo Fisher Scientific) assays for quantity and FEMTO Pulse (Agilent) for quality. For HiFi sequencing, samples exhibiting a mode size above 50Lkbp were considered to be good candidates. Libraries were prepared using the SMRTbell Express Template Prep Kit 2.0 (PacBio). In brief, 12 μl of DNA was first sheared using gTUBEs (Covaris) to target 15–18Lkbp fragments. Two 5 µg of sheared DNA were used for each prep. DNA was treated to remove single-stranded overhangs, followed by DNA damage repair and end repair/A-tailing. The DNA was then ligated with a V3 adapter and purified using Ampure beads. The adapter ligated library was treated with Enzyme mix 2.0 for nuclease treatment to remove damaged or non-intact SMRTbell templates, followed by size selection using Pippin HT (Sage Science) generating a library with a size >10Lkbp. The size-selected and purified >10Lkbp fraction of libraries was used for sequencing on the Sequel II (PacBio) system.

### Genome assembly and SD annotation

PacBio HiFi sequencing data from 1KG samples were assembled using hifiasm ^29^. Because of the difficulties in mapping acrocentric SDs, we excluded sequence mapping to the short arms of chromosomes 13, 14, 15, 21 and 22. The autosomal contigs are scaffolded using RagTag (v2.1) ^70^. We masked repeat content using RepeatMasker (v4.1) and called SDs using SEDEF (>90% and >1 kbp) ^71^. We matched Illumina short-read sequencing data for all 170 haplotypes, which were used for additional read-depth support of the putative duplicated regions (fastCN) ^72^.

### Variant calling

We used PAV, an assembly-based phased assembly variant caller, to call variants for 164 genome assemblies, 58 of which were phased into paternal and maternal haplotypes using parental Illumina short-read data (https://github.com/EichlerLab/pav; v.2.1.0). The regions that align 1-to-1 via minimap2 ^73^ “-x asm20 --secondary=no -s 25000 -K 8G”, showing no variants, were assigned as the reference, 0|0 genotype, while the regions outside of the alignment blocks were considered as missing genotypes when merging the variant calls of individual samples. This was done by via BCFtools merge --missing-to-ref followed by BCFtools view with the aligned regions (v.1.9) ^74^. In addition, in order to focus on confident variant call set, we additionally defined a 1-to-1 alignment block, which is syntenic with length >1 Mbp, in at least 80% of the samples. The population statistics were calculated across this 1- to-1 syntenic, shared alignment blocks of length 2.605 Gbp (90%), across the autosomes.

### Iso-Seq and transcript analyses

We used long-read RNA-seq (PacBio Iso-Seq) data to look for evidence of expression of newly discovered low-frequency gene duplications. Examining regions where 10 or fewer haplotypes have a duplication relative to the remainder of the samples, we align 563 million FLNC reads from 241 libraries to *de novo* assemblies and T2T-CHM13 v2.0 as a reference. Only alignments with >99.9% identity to the novel duplication and <99.7% identity to T2T-CHM13 were considered. To generate gene models for each *de novo* assembly, we transferred GENCODE v44 gene models with Liftoff (v1.6.3) ^75^, classified Iso-Seq reads compared to the Liftoff gene models with PacBio Pigeon and SQANTI3 (v5.2) ^76^, and predicted open reading frame sequences with GeneMark ^77^. Each coding gene prediction was compared to the NCBI nonredundant protein database (nr) with BLAST (v.2.15) ^78^. We limited our Iso-Seq analysis to transcripts aligning to reference SDs (227.4 Mbp), their boundaries (defined as 10% of the SD block size on each edge, 28.1 Mbp), SDs seen in at least one assembled haplotype but not the reference (17.9 Mbp), and regions corresponding to highly divergent loci in the *de novo* assemblies (unaligned to T2T-CHM13 with the asm20 preset of minimap2 but forced to align with -r2k,200k -N50 parameters, 202.7 Mbp), totaling 476.1 Mbp of theT2T-CHM13 genome. Gene ontology was performed with g:Profiler (database e111_eg58_p18_30541362) ^79^.

### Copy number estimation

#### Assembly-based methods

To estimate copy number, we mapped protein-coding genes (genic sequences from T2T-CHM13) overlapping with SDs to each haplotype assembly using minimap2 (>60% coverage and >90% identity)^73^. Single exons or short genes (coding sequence < 200 bp) were excluded. If genes are composed with high repeat content, copy number can be overestimated due to partial mapping of repeat content. To remove this incorrect mapping, we removed alignments that completely matched the repeat sequence. To increase contiguity of the alignment and to estimate counts of high copy number genes, we customized the minimap2 options as follows: ‘minimap2 -cx map-ont -f 5000 -k15 -w10 -p 0.05 -N 200 -m200 - s200 -z10000 --secondary=yes --eqx ’. To avoid any bias due to switch errors, we estimated gene copy number in diploid genomes. We also excluded duplicate genes found only in one haplotype.

#### Illumina fastCN

Read-depth copy number was estimated with fastCN ^72^, using Illumina reads for each sample and comparing to the T2T-CHM13 genome. To remove signal from variable number tandem repeats (VNTRs), we excluded fastCN windows that overlapped TRF ^80^ or WindowMasker ^81^ calls by more than 10%; to estimate gene copy number, we only considered windows contained within the bounds of each annotated gene, taking the median value. To check for biased copy number estimates, we decomposed the T2T-CHM13 genome into 36-mers and estimated copy number with the fastCN pipeline, simulating fastCN results for a perfectly matched sample and reference. By default, fastCN overcorrected read depth based on GC content for these unbiased artificial reads, due to overrepresentation of extreme GC values in the human genome itself. We recalibrated the GC correction, window-by-window, based on these results. Read-depth-based methods also underestimate copy number to a variable extent based on sequence divergence between paralogs, due to unaligned sequence. To correct for this, we calculated an adjustment factor for each gene to match fastCN results to alignment-based assembly copy number with T2T-CHM13 as ground truth. Genes that required more than a 50% adjustment to match copy number between methods were excluded from further analysis (n=82).

## DATA AVAILABILITY

The raw sequencing data generated in this study are available under project ID PRJEB58376 (https://www.ebi.ac.uk/ena/browser/view/PRJEB58376) and the HPRC year 1 PacBio HiFi data is available under PRJNA730823 (https://ncbi.nlm.nih.gov/bioproject/PRJNA730823) or https://github.com/human-pangenomics/HPP_Year1_Assemblies. The raw genome sequencing data generated from this study are available online (https://ftp.1000genomes.ebi.ac.uk/vol1/ftp/data_collections/HGSVC3/).

## COMPETING INTERESTS

E.E.E. is a scientific advisory board (SAB) member of Variant Bio, Inc. C.L. is an SAB member of Nabsys and Genome Insight. The other authors declare no competing interests.

## ACKNOWLEDGMENTS

We thank T. Brown for assistance with manuscript editing and preparation. This research was supported, in part, by funding from the National Institutes of Health (NIH) grants R01 HG002385, R01 HG010169, and U24 HG007497 (to E.E.E. and C.L.). E.E.E. is an investigator of the Howard Hughes Medical Institute.

This article is subject to HHMI’s Open Access to Publications policy. HHMI lab heads have previously granted a nonexclusive CC BY 4.0 license to the public and a sublicensable license to HHMI in their research articles. Pursuant to those licenses, the author-accepted manuscript of this article can be made freely available under a CC BY 4.0 license immediately upon publication.

