## Supplementary figures for "Structural polymorphism and diversity of human segmental duplications"

**SUPPLEMENTARY FIGURES for Jeong et al.**

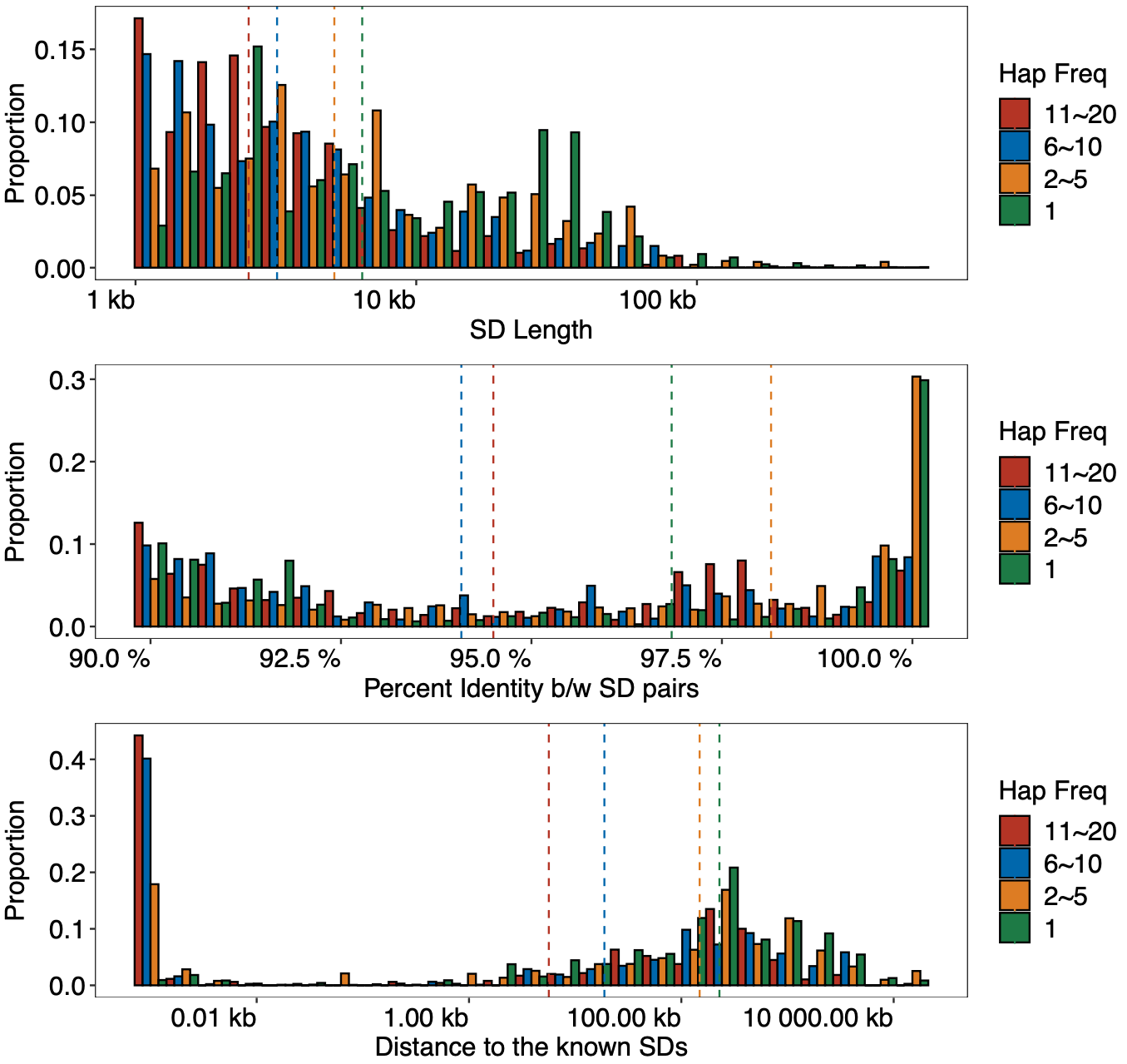

**Supplementary Figure 1. Histogram comparing the sequence identity and length of polymorphic segmental duplications (SDs) at different haplotype frequencies.**

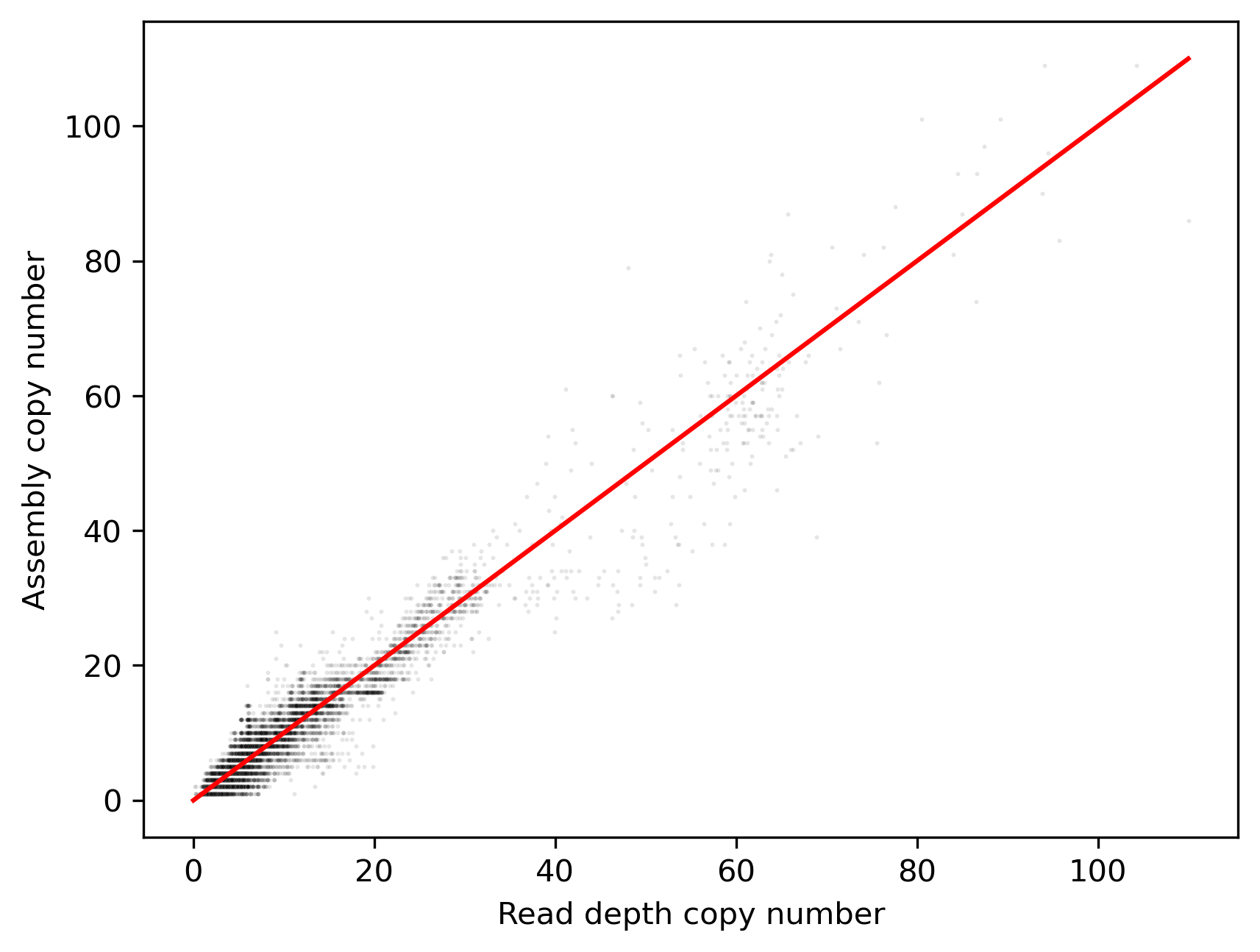

**Supplementary Figure 2. Read-depth-based copy numbers estimated with fastCN compared to assembled copy number for each sample, summed between the two haplotypes.** Each point represents the copy number estimates for a gene family in a sample (R2 = 0.94).

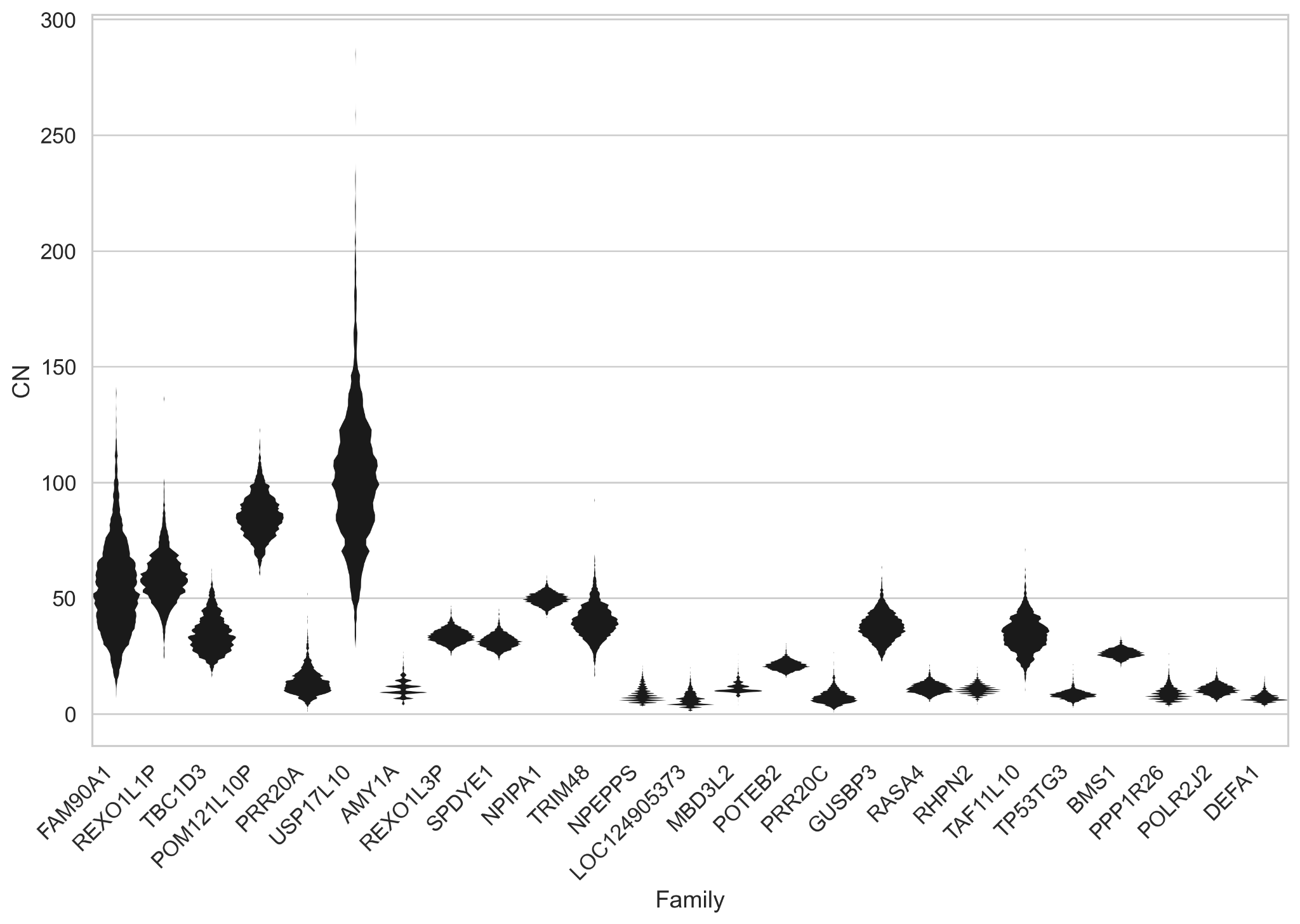

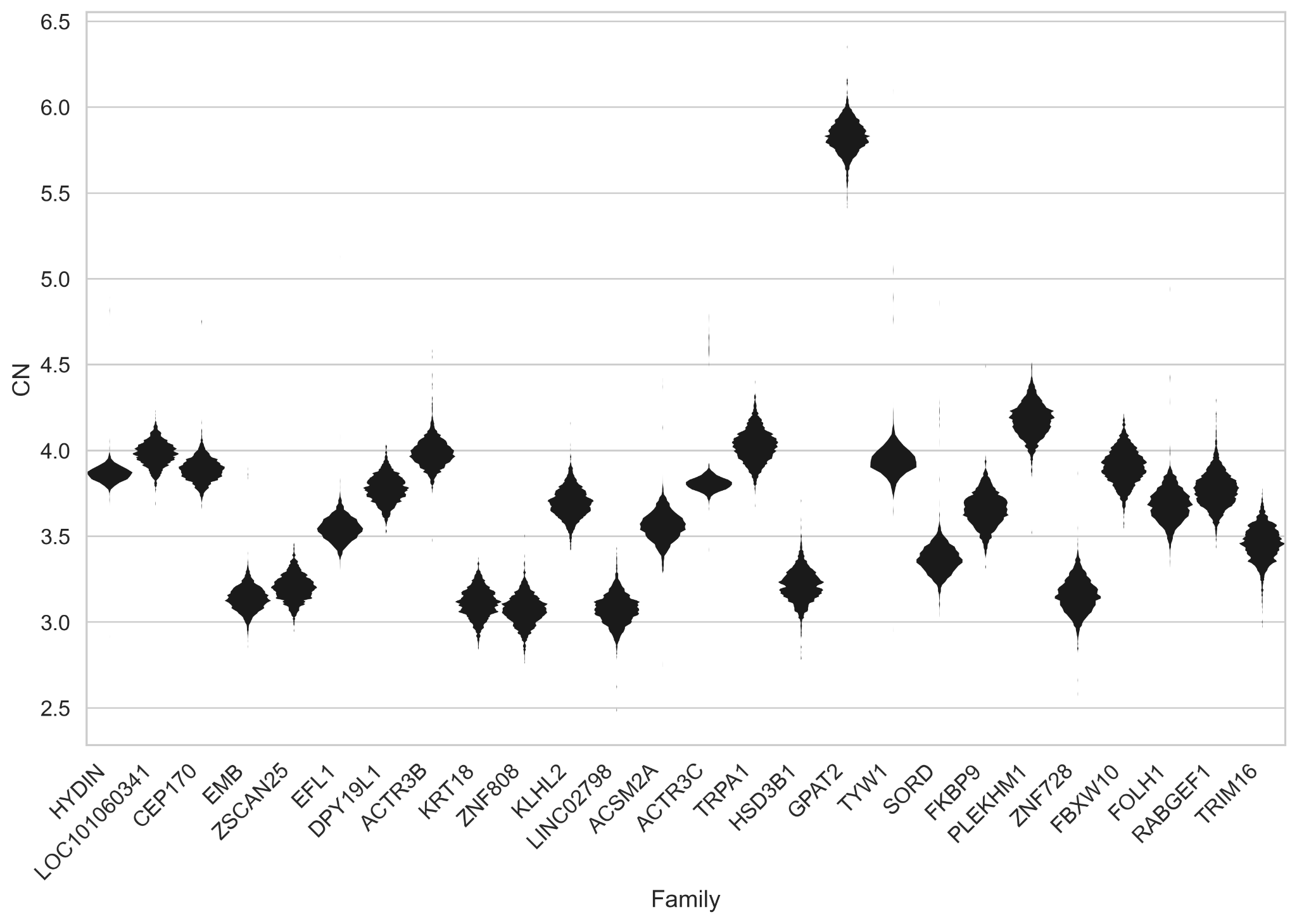

**Supplementary Figure 3. Copy number distribution of high- and low-variance gene families.** The read-depth copy number of gene families with highly variable (above) and nearly fixed copy number (below) are displayed. Gene families are selected and ordered by variance, requiring an average diploid copy number greater than three.

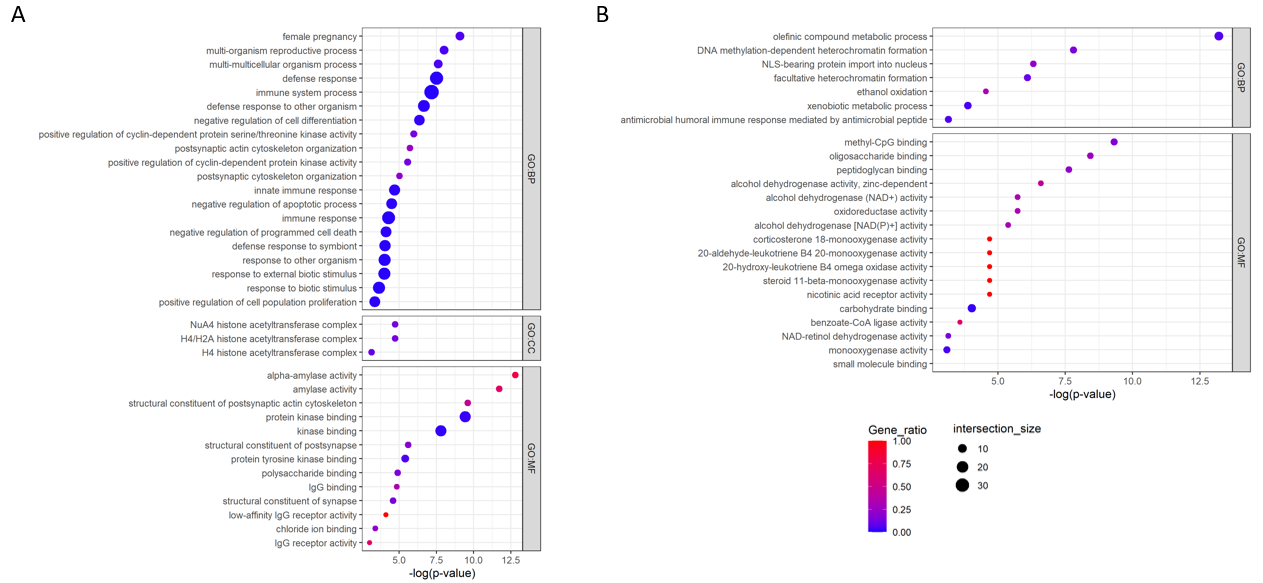

**Supplementary Figure 4. Gene ontology enrichment of the (A) top 100 variable gene families (n = 358) and (B) invariable genes (n = 115).** X-axis indicates negative log transformed adjusted p-value. The number of intersecting genes is indicated by the size of the circle and the gene ratio represents the number of intersect/term size.

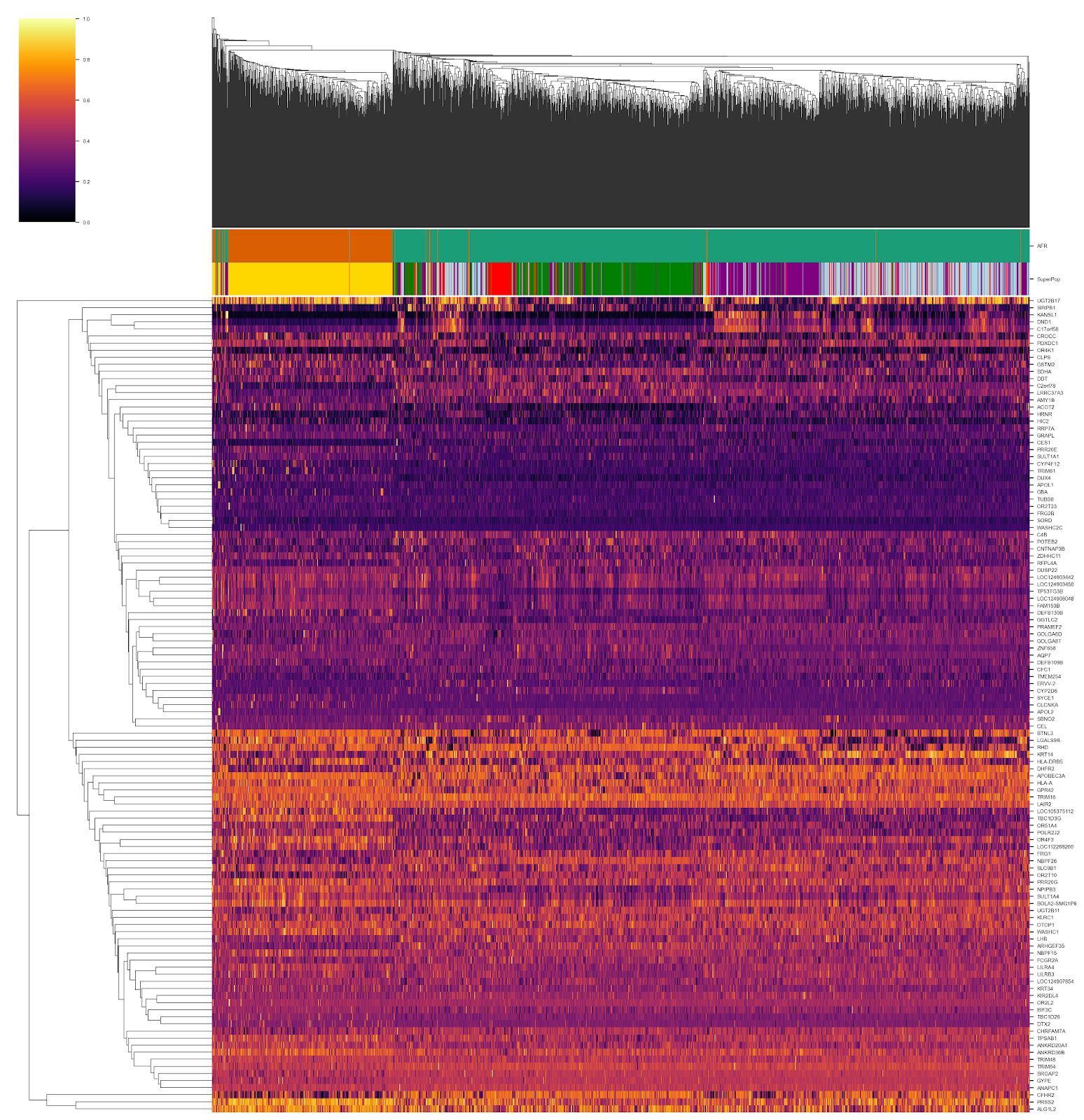

**Supplementary Figure 5. Population-stratified genic copy number in 2,196 unrelated individuals from the 1000 Genomes Project.** Gene copy number values are centered on the mean for each gene and scaled by unit variance to range from 0-1. One paralog per gene family and duplication block is shown. 73/115 deduplicated population-stratified genes have higher mean copy number in the African group as compared to the non-African group (p=0.002). Superpopulations as described in the 1000 Genomes Project are shown above (Africa: gold, East Asia: green, South Asia: purple, Europe: blue, the Americas: red). Copy number estimates by population group (below).

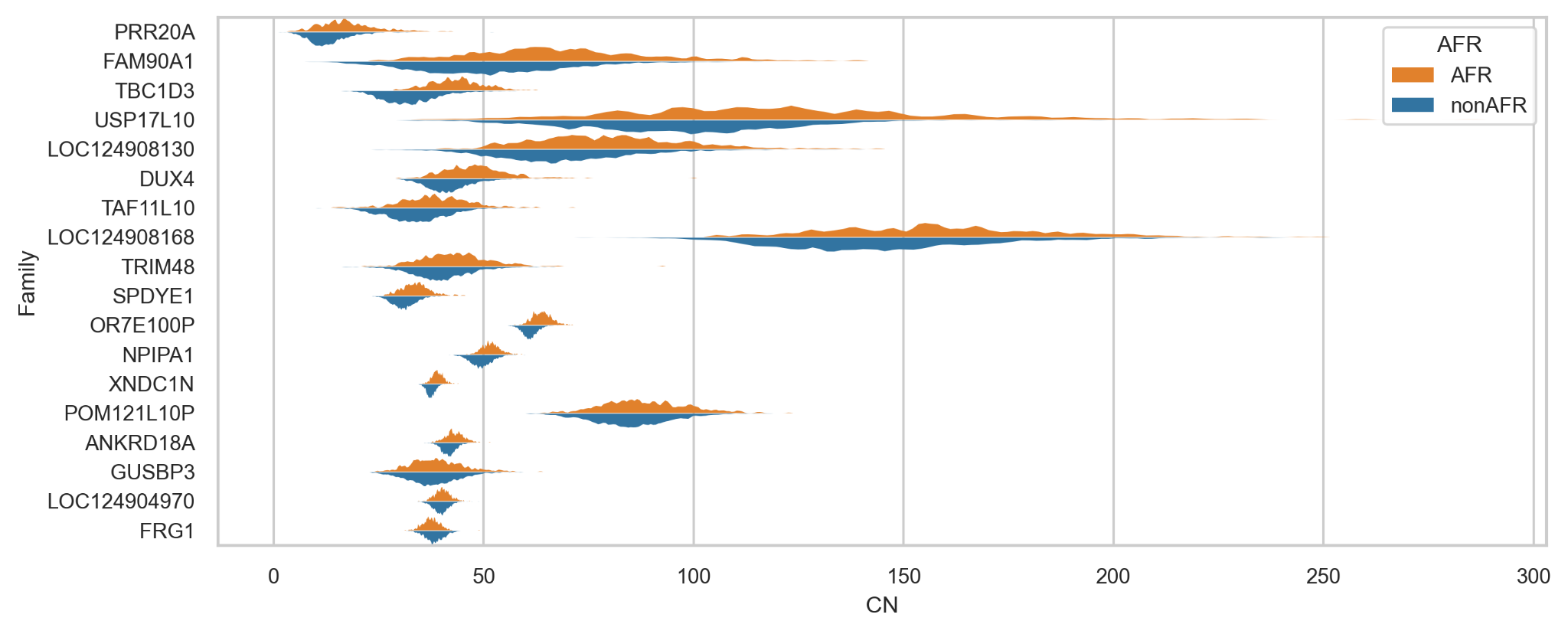

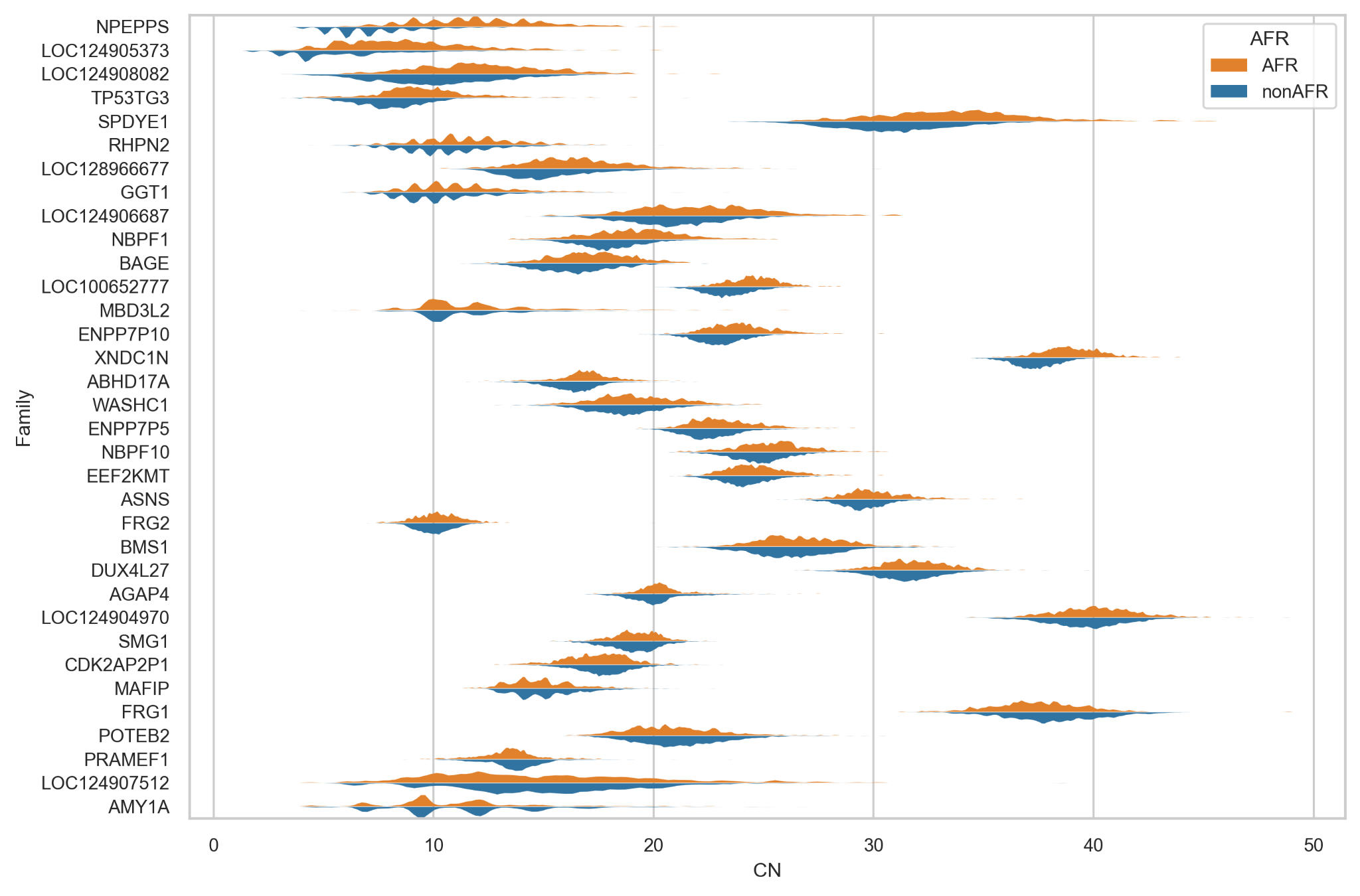

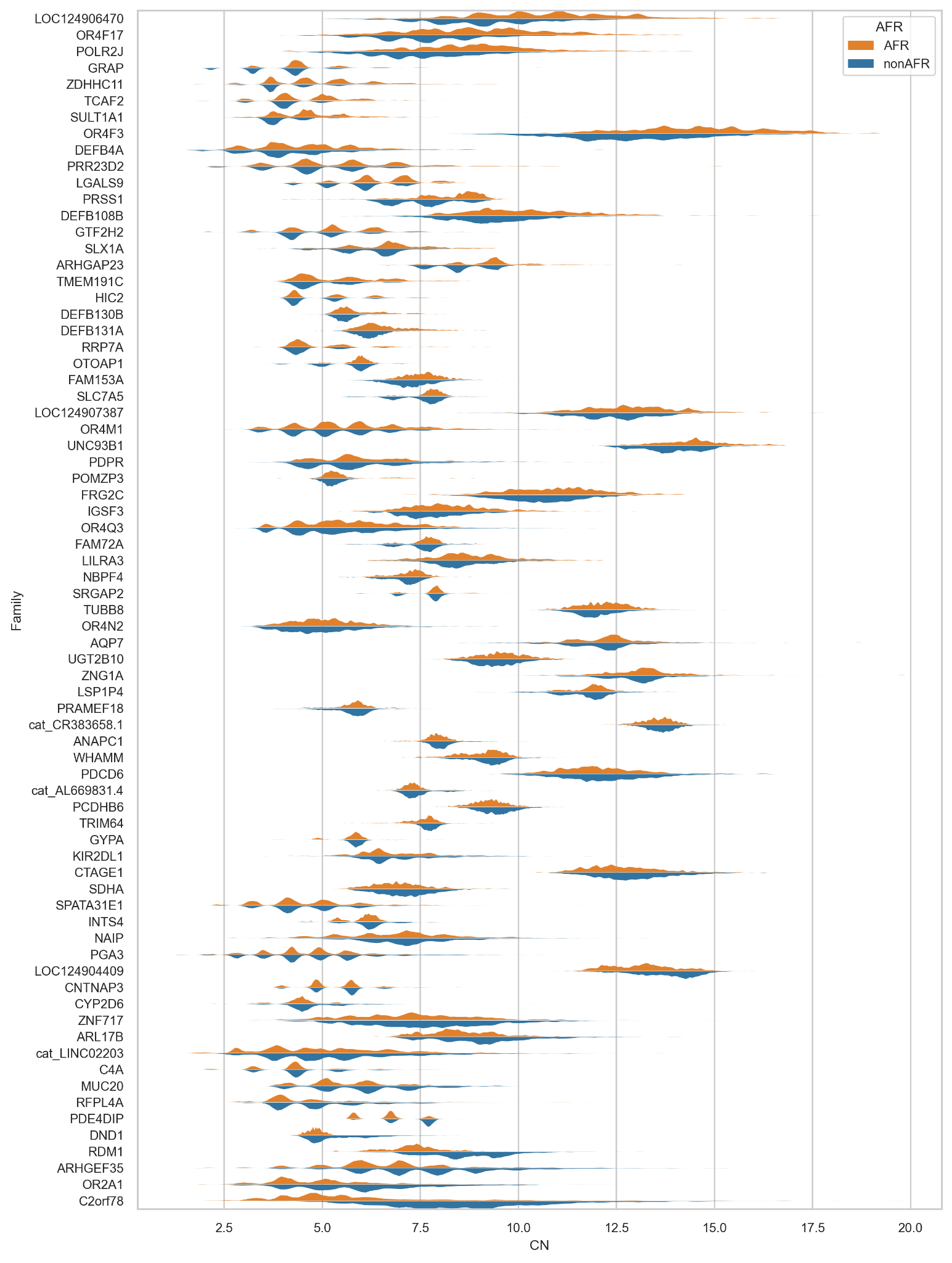

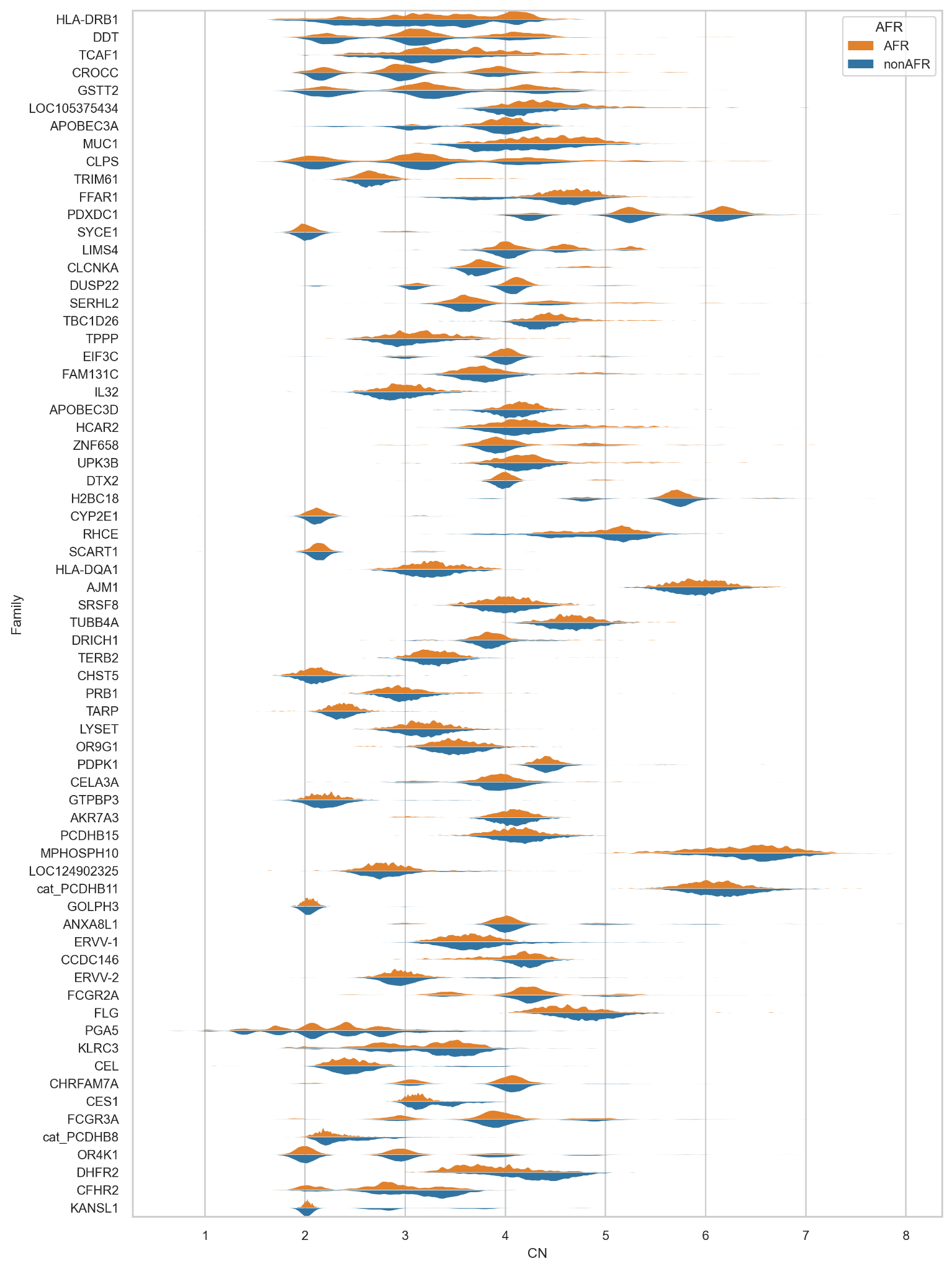

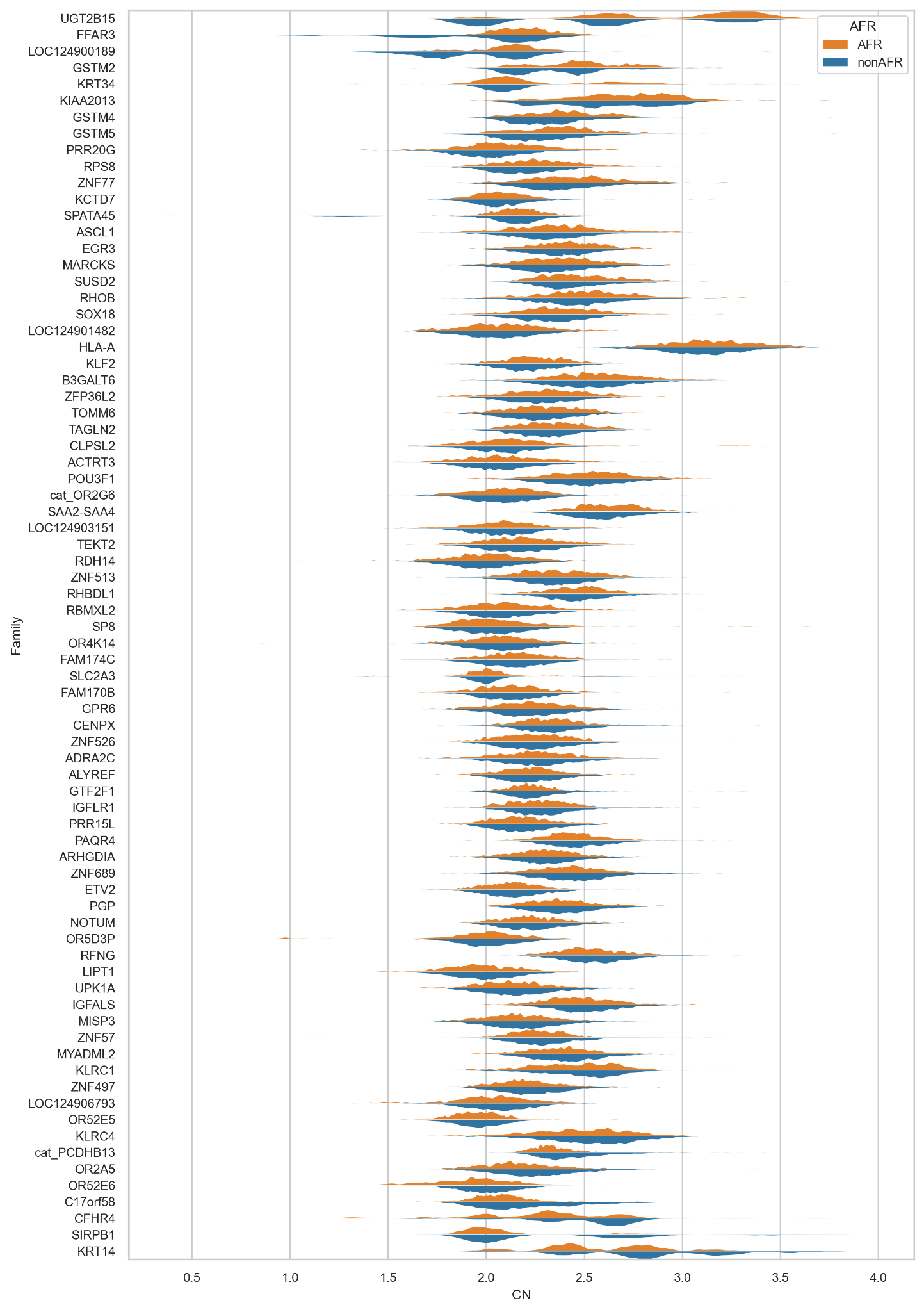

**Supplementary Figure 6. Population-stratified gene copy number in 1000 Genomes Project.** To validate population differences in gene copy number, we repeated the analysis on an unrelated subset of high-coverage Illumina data from the 1000 Genomes Project, excluding samples identified as related by published pedigree or with cryptic relationships (2nd degree or less) as identified by Somalier (Pedersen, Genome Medicine 2020). Individuals from the highly-admixed ACB, ASW, and PUR populations were also excluded, leaving n=2,196.

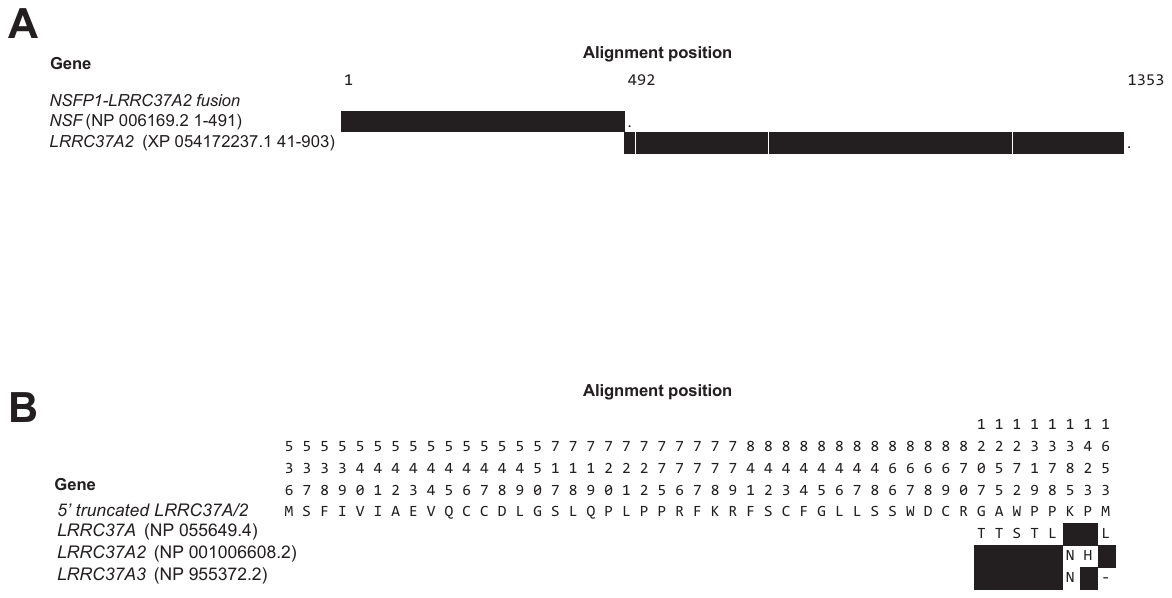

**Supplementary Figure 7. Multiple sequence alignment of novel *LRRC37A* family gene predictions compared to reference annotations.** Black boxes indicate matches between the novel prediction and reference annotations. (A) The *NSFP1-LRRC37A2* gene prediction encodes a fusion between *NSFP1* and *LRRC37A2*. (B) The 5' truncated *LRRC37A/2* gene prediction has 39 novel amino acids followed by a hybrid of *LRRC37A* and *LRRC37A2*.
